# Genotype-dependent transcriptional trajectories during prolonged heat stress in *Capsicum annuum* L

**DOI:** 10.64898/2026.08.03.742462

**Authors:** Matteo Martina, Edoardo Vergnano, Francesca Secchi, Anna Maria Milani, Lorenzo Barchi, Andrea Moglia, Alberto Acquadro, Cinzia Comino, Ezio Portis

## Abstract

Heat stress is one of the most damaging abiotic constraints on crop productivity, and its consequences are expected to intensify as extreme temperature events become more frequent and severe. Pepper (*Capsicum annuum* L.) is particularly vulnerable to sustained high temperatures, which can disrupt photosynthetic performance, cellular homeostasis, and redox regulation. However, the physiological and transcriptional dynamics underlying genotype-dependent responses to prolonged heat exposure remain insufficiently understood. We combined repeated physiological measurements with time-course RNA sequencing to compare GPC003240, previously identified as a candidate heat-tolerant accession, with two non-elite accessions, GPC010350 and GPC014930, which are phenotypically divergent from each other, under 40/30 °C Day/night temperatures for up to six days. GPC010350 maintained comparatively stable photosystem II performance and higher stomatal conductance, whereas GPC014930 showed progressive photochemical impairment and lower conductance; GPC003240 displayed a distinct, moderately responsive profile. Transcriptomic responses showed partial functional convergence during the early phase of stress exposure but diverged markedly after six days. When gene expression at day 6 was compared with the pre-treatment baseline separately within each genotype, 4,436 differentially expressed genes were detected in GPC010350, compared with 680 in GPC003240 and only 78 in GPC014930. The late response of GPC010350 was associated with enrichment of RNA- and ribosome-related, biosynthetic, DNA-repair, and genome-maintenance functions. By contrast, GPC014930 showed negative enrichment of photosynthesis, plastid organization, redox homeostasis, and translation-related processes. Global co-expression analysis identified a time-decreasing photosynthesis-associated module (ME5) and two time-increasing modules, ME12 and ME19, that were enriched in genes contributing to the late GPC010350 response. Integration of differential expressions, module membership, and functional annotation highlighted a heat shock transcription factor (*Caz03g27980*), *HSP101* (*Caz03g07770*), and a dual-specificity phosphatase (*Caz05g20970*) as candidates for further investigation. Overall, the results suggest that genotype-dependent responses to prolonged heat exposure were associated not only with the magnitude of early transcriptional change, but also with differences in the temporal organization of stress-response, maintenance, and metabolic processes. The contrasting responses of the non-elite accessions GPC010350 and GPC014930 further highlight the value of phenotypically diverse germplasm for uncovering mechanisms relevant to future heat-tolerance breeding.

## 1 Introduction

Heat stress is a major constraint on crop productivity, and its effects are expected to become increasingly severe as extreme temperature events intensify in frequency and duration. Bell pepper (*Capsicum annuum* L.) is a crop of substantial economic and nutritional value, yet it is highly susceptible to elevated temperatures, particularly during reproductive development, when heat can markedly reduce fruit set, yield, and quality. Although pepper generally performs optimally between 18 and 30 °C, temperatures above 35 °C can disrupt cellular homeostasis by promoting protein denaturation, membrane destabilization, photosynthetic inhibition, and excessive production of reactive oxygen species (ROS - Guo et al., 2014; Tang et al., 2022; Kim et al., 2023).

Plant acclimation to high temperature depends on extensive and multilayered transcriptional reprogramming. Canonical heat-response pathways are coordinated by heat shock factors (HSFs) and heat shock proteins (HSPs), but sustained adaptation also requires broader adjustments in hormone signaling, redox regulation, primary and secondary metabolism, RNA processing, protein synthesis, and organellar function (Liu et al., 2011; Darriere et al., 2022a; Zheng et al., 2023). These mechanisms have been investigated extensively in model plants and major crops, including *Arabidopsis*, tomato, and rice (Liu et al., 2011; Kostaki et al., 2020; Vitoriano and Calixto, 2021). In pepper, however, transcriptome-wide studies remain comparatively limited, particularly those involving genetically and phenotypically diverse accessions and time-course designs capable of distinguishing early responses from transcriptional programmes maintained or emerging during prolonged stress (Li et al., 2015; Wang et al., 2019, 2021; Tang et al., 2022).

A central challenge in stress transcriptomics is to distinguish broadly conserved heat-response mechanisms from genotype-dependent programmes. This distinction is particularly relevant for crop improvement, because contrasting stress performance may depend not only on the activation of an initial defence response, but also on the capacity to sustain and reorganize cellular functions during prolonged exposure. Moreover, phenotypically divergent and non-elite germplasm may harbour response mechanisms that are poorly represented in commercial material but could provide useful sources of variation for future breeding. Time-course RNA sequencing, combined with factorial models accounting for genotype, exposure duration, and their interaction, offers a suitable framework for resolving shared and divergent components of the heat-associated transcriptome.

To investigate these processes, we selected three *C. annuum* accessions from the G2P-SOL core collection using complementary phenotypic criteria (Tripodi et al., 2021). GPC003240 was included as a candidate heat-tolerant accession because it retained a control-like phenotypic profile under heat stress in previous phenomics-based screening (Fumia et al., 2023). By contrast, no prior heat-stress phenotyping was available for GPC010350 and GPC014930; these two non-elite accessions were instead selected for their broader phenotypic divergence and genetic distinctness from commercial pepper ideotypes, and their heat-stress responses were characterized here for the first time. This design was intended to evaluate contrasting responses to prolonged heat exposure and to explore whether commercially underutilized germplasm could reveal physiological and molecular mechanisms of potential relevance for heat-tolerance breeding. Plants were exposed to a continuous 40/30 °C Day/night regime and sampled for RNA-seq at the common pre-treatment baseline and after two (T2), three (T3), and six (T6) days of exposure, with three biological replicates per genotype and time point. Physiological measurements were collected at 24-h intervals throughout the six-day exposure period to characterize exposure-associated responses and define condition-specific trajectories. RNA-seq data were analyzed using a full factorial *DESeq2* model to partition genotype, exposure-time, and genotype-by-time effects. Functional interpretation was performed through Gene Ontology-based gene set enrichment analysis, while weighted gene co-expression network analysis was used to identify exposure-time-associated and genotype-associated transcriptional programmes. By integrating physiological and transcriptomic evidence, this study aimed to resolve shared and genotype-dependent responses to prolonged heat exposure and to identify candidate processes, modules, and genes with potential relevance for future pepper improvement.

## 2 Materials and Methods

### Plant material and heat-stress treatment

Three *Capsicum annuum* L. genotypes from the G2P-SOL core collection were included in the experiment: GPC003240, GPC010350, and GPC014930 (Tripodi et al., 2021; G2P-SOL, https://www.g2p-sol.eu/). GPC003240 was selected as a candidate heat-tolerant accession based on the phenomics-based screening reported by Fumia et al. (2023). GPC010350 and GPC014930, for which no prior heat-stress phenotyping was available, were selected as non-elite accessions with broader phenotypic divergence from each other, to extend the range of responses examined beyond GPC003240’s known profile. Any reference in this study to comparatively tolerant, sensitive, or intermediate behavior denotes the physiological response observed under the present experimental regime and does not represent a priori classification. Four plants per genotype were grown to the vegetative stage over a two-month period in pots containing approximately 300 g of soil, under controlled conditions in an Aralab FITOCLIMA 600 growth chamber (Aralab, Portugal). Before heat treatment, plants were maintained under a 12 h photoperiod, 350 µmol photons m⁻² s⁻¹ photosynthetically active radiation, 30/25 °C Day/night temperature, and 70–80% relative humidity. This humidity range was selected to provide moderately humid, non-water-limiting atmospheric conditions and is consistent with conditions commonly used for *Capsicum* (Dang et al., 2018; Rajametov et al., 2021; Wang et al., 2021). The three genotypes were randomized among chamber positions according to a randomized complete block design, and plants were periodically rotated to minimize positional effects. Uniform irrigation and fertilization were maintained throughout the growth and treatment periods. Heat exposure was initiated immediately after the common pre-treatment baseline measurement, T0, by increasing the chamber setpoints to 40/30 °C Day/night while maintaining the same photoperiod, irradiance, and relative-humidity range. The treatment was applied continuously for six days.

### Physiological measurements

Physiological measurements were performed in the morning at T0, immediately before the initiation of heat treatment, and subsequently at 24-h intervals from T1 to T6, corresponding to one to six days of heat exposure. This measurement schedule was denser than the RNA-seq sampling time course (T0, T2, T3, and T6), allowing physiological trajectories to be resolved at a finer temporal resolution than the transcriptomic dataset. The measurement schedule therefore included all RNA-seq sampling time points. To minimize disturbance of the chamber environment, plants were temporarily transferred to the adjacent laboratory for physiological measurements. Before measurement, plants were allowed to equilibrate for 10 min under standardized laboratory conditions. The same equilibration period and measurement procedures were applied consistently across genotypes and sampling days.

Chlorophyll fluorescence parameters, including the maximum efficiency of photosystem II in the light-adapted state (Fv′/Fm′), the effective quantum yield of photosystem II photochemistry (ΦPSII), and non-photochemical quenching (NPQt), were measured using a MultispeQ V2.0 device (PhotosynQ). All four plants per genotype were measured repeatedly with the MultispeQ device throughout the time course. Three of these four plants were additionally used for LI-600 gsw measurements, Ψleaf assessment, and RNA-seq leaf sampling; the fourth plant was measured only with the MultispeQ device and was not involved in any other measurement. Leaf-level measurements were averaged to obtain one biological value per plant. Incident photosynthetically active radiation recorded during each measurement was retained to characterize the laboratory measurement environment.

Stomatal conductance to water vapour (gsw) and transpiration rate were measured after the standardized 10-min equilibration period using an LI-600 Porometer/Fluorometer (LI-COR Biosciences, Lincoln, NE, USA). For each genotype and time point, measurements were collected from two fully expanded, visibly dry leaves on each of the same three plants used throughout the study. Raw observations were subjected to quality control, after which stomatal conductance and transpiration were recalculated at the individual-leaf level using the psychrometric temperature correction described by Rizzo and Bailey (2026), which accounts for temperature changes along the instrument airflow path. A stomatal-sidedness factor of 1.0 was used; corrected conductance values therefore refer to the leaf surface directly measured by the instrument. The two corrected leaf-level observations were subsequently averaged to obtain one biological value per plant. Measurement time, incident PAR, inlet humidity and vapour pressure, and instrument temperature variables recorded by the LI-600 were retained for quality control and characterization of the laboratory measurement conditions. Leaf and instrument temperature variables were used for implementation of the psychrometric correction but were not analyzed as physiological response traits.

Leaf water potential (Ψleaf) was measured only at T0 and T6, rather than at every physiological time point, because pressure-chamber measurement is destructive to the sampled leaf; intermediate time points were not sampled because plants did not yet display visible signs of stress, and Ψleaf was therefore assessed at the pre-treatment baseline (T0) and at the time point when visible stress symptoms became apparent (T6), on fully expanded, transpiring leaves using a Scholander-type pressure chamber (Model 1505D; PMS Instrument Co., Albany, OR, USA), following Secchi et al. (2012). For each genotype and time point, one leaf at the same physiological state was collected from each of the same three plants used for RNA-seq sampling and gsw measurements.

Repeatedly measured physiological traits were analyzed in R using linear mixed-effects models, with genotype, time point, and their interaction included as fixed effects and plant identity included as a random effect to account for repeated measurements of the same individuals. Leaf water potential was analyzed using a factorial model including genotype, time point, and their interaction. When the genotype × time interaction was significant, estimated marginal means were compared among genotypes within each time point using pairwise contrasts with Holm adjustment for multiple testing. Statistical significance was assessed at α = 0.05. Error bars represent standard deviations unless otherwise stated.

### RNA extraction, library preparation, and sequencing

For transcriptome profiling, fully expanded leaves were collected at T0, immediately before the initiation of heat treatment, and after two, three, and six days of exposure (T2, T3, and T6, respectively). For each genotype and time point, three biological replicates were sampled, each consisting of leaf tissue collected from an individual plant. Samples were immediately frozen in liquid nitrogen and stored at −80 °C until processing. Total RNA was extracted from approximately 100 mg of powdered leaf tissue using the Spectrum™ Plant Total RNA Kit (Sigma-Aldrich), following the manufacturer’s protocol. RNA concentration and purity were assessed by NanoDrop spectrophotometry and Qubit fluorometry, whereas RNA integrity was evaluated by agarose gel electrophoresis and capillary electrophoresis, as depicted in Martina et al. (2026). Stranded mRNA libraries were prepared from poly(A)-enriched RNA according to standard Illumina protocols and sequenced by IGAtech (Udine, Italy) on an Illumina NovaSeq X platform, generating 2 × 150-bp paired-end reads. Sequencing yielded approximately 40 million reads per sample. Raw sequencing data have been deposited in the NCBI Sequence Read Archive under BioProject accession SUB16378501.

### Read quantification and gene-level summarization

Raw sequencing reads were trimmed and filtered using fastp (Chen et al., 2018) with standard parameters. Transcript-level abundances were estimated with Salmon (Patro et al., 2017) in quasi-mapping mode against the *C. annuum* cv. Zhangshugang reference transcriptome (Liu et al., 2023), using the --validateMappings and --gcBias options. Transcript-level estimates were summarized to gene-level counts using the tximport package (Soneson et al., 2015) in R (R Core Team, 2020), with a transcript-to-gene mapping derived from the reference genome annotation. No manual low-count pre-filter was applied; all genes were retained in the DESeq2 model, and low-count genes were excluded from multiple-testing correction via DESeq2’s default independent filtering.

### Differential expression analysis

Differential expression analysis was performed using *DESeq2* (Love et al., 2014). A full interaction model was specified as ∼ genotype + time + genotype:time, with GPC003240 set as the reference genotype level and T0 as the reference time point. Size factor estimation and negative binomial model fitting were performed using the DESeq function with default settings. Log2 fold changes were shrunk using the *lfcShrink* function (type = “ashr”) to produce more stable effect-size estimates for genes with low counts or high dispersion; adjusted p-values are unaffected and correspond to the original Wald test. Variance-stabilized expression values were obtained using the VST transformation (blind = FALSE) and used for exploratory analyses, including principal component analysis and sample-to-sample Euclidean distance heatmaps. Pre-specified contrasts were extracted to evaluate three classes of transcriptional differences: temporal responses within each genotype relative to T0, obtained by comparing each heat-exposure time point with T0; pairwise genotype differences at each time point; and genotype-by-time interaction terms, representing differences in exposure-associated temporal log2 fold changes between GPC010350 or GPC014930 and the reference genotype GPC003240. A complete list of contrasts, DESeq2 coefficient names or contrast vectors, and DEG counts per contrast is provided in Supplementary Table S1. Genes were classified as differentially expressed when they met a Benjamini–Hochberg-adjusted p-value < 0.05 and an absolute log2 fold-change > 0.5. DEG counts were interpreted as estimates of the breadth of detectable transcriptional reprogramming rather than as exhaustive inventories of biologically validated heat-response genes. To assess the robustness of the main patterns to threshold choice, a sensitivity analysis reporting DEG counts under alternative log2 fold-change cutoffs is provided in Supplementary Table S2. Key biological conclusions were further evaluated using ranked Gene Set Enrichment Analysis (GSEA), which does not rely on binary DEG classification. Set-theoretic comparisons among genotype-specific heat-responsive DEG sets were performed at each time point using UpSet plots implemented with the *ComplexUpset* package (Lex et al., 2014; Michał Krassowski et al., 2022) and Venn diagrams generated with *ggvenn* (Yan, 2021).

### Gene Set Enrichment Analysis

Functional enrichment analysis was performed using gene set enrichment analysis (GSEA) implemented with the *gseGO* function of the *clusterProfiler* package (Yu et al., 2012). A custom Gene Ontology annotation database was generated from functional annotations obtained with eggNOG-mapper (Cantalapiedra et al., 2021) applied to the *C. annuum* cv. Zhangshugang protein sequences. Gene identifiers in the DESeq2 results were verified to match the *OrgDb* key type used for enrichment analysis. For each contrast, genes were ranked by the *DESeq2* Wald statistic, which jointly reflects estimated effect size and precision. GSEA was performed separately for the Gene Ontology Biological Process (BP), Molecular Function (MF), and Cellular Component (CC) categories, using minimum and maximum gene-set sizes of 10 and 500 genes, respectively. Enrichment direction and magnitude were summarized using the normalized enrichment score (NES). Positive NES values indicate enrichment toward genes with positive Wald statistics, whereas negative NES values indicate enrichment toward genes with negative Wald statistics. For within-genotype contrasts, these directions correspond to comparatively higher or lower expression at the indicated exposure time relative to T0. For genotype-by-time interaction contrasts, they indicate comparatively more positive or more negative temporal responses in GPC010350 or GPC014930 relative to GPC003240. Statistical significance was assessed using a Benjamini–Hochberg-adjusted *p*-value threshold of 0.10, without applying an additional NES cutoff. The *nPermSimple* parameter was set to 20,000. GSEA was conducted for within-genotype exposure contrasts at T2 and T6 and for the genotype-by-time interaction contrasts at T6 comparing GPC010350 and GPC014930 with GPC003240. Results were visualized using ridge plots, dot plots, and enrichment-score plots for representative gene sets.

### Weighted Gene Co-expression Network Analysis

Co-expression network analysis was performed in R using the WGCNA package (Langfelder and Horvath, 2008). Two complementary analyses were conducted: a global network including all 36 samples (3 genotypes x 4 timepoints x 3 replicates) and exploratory per-genotype networks. For the global network, VST-normalized expression values from all samples were used, and the 50% most variable genes, based on sample variance, were retained. Sample quality was assessed using the *goodSamplesGenes* function. A signed co-expression network was constructed using the *blockwiseModules* function. The soft-thresholding power was selected to achieve a scale-free topology fit index (R²) > 0.85. Network construction used a minimum module size of 30 genes, a merge cut height of 0.25, and a maximum block size of 20,000 genes. Module eigengenes were computed, and modules with similar eigengene profiles were merged according to the specified cut height. Module–trait correlations were calculated between module eigengenes and trait vectors representing heat exposure time and genotype identity. Heat exposure time was coded in hours as 0, 48, 72, and 144, corresponding to T0, T2, T3, and T6, respectively. Genotype identity was encoded using binary indicator variables. Pearson correlation was used to estimate module–trait associations, and significance was assessed using the Student’s t-distribution approximation. A summary of all modules, including size, module–trait correlations, top hub genes, and enriched GO terms, is provided in Supplementary Table S3. Exploratory per-genotype networks were constructed separately for each genotype using 12 samples per genotype, corresponding to four time points and three biological replicates. For these analyses, the 40% most variable genes were retained and a minimum module size of 15 genes was used. Because these networks were inferred from a limited number of samples, they were interpreted as exploratory and used to nominate candidate co-expression programs rather than to define definitive genotype-specific regulatory structures. Module–trait correlations were computed against heat exposure time. The significance of module eigengene trends over time was assessed by linear regression. Module overlap between per-genotype networks was evaluated using pairwise hypergeometric tests, followed by Benjamini–Hochberg correction. Module preservation was assessed using the WGCNA modulePreservation function with 100 permutations, using GPC003240 as the reference network.

### Statistical analysis and data visualization

All statistical analyses were performed in R (R Core Team, 2020). Principal component analysis was conducted using the *prcomp* function on VST-normalized expression values. Figures were generated using *ggplot2* (Wickham, 2016), *pheatmap* (Kolde, 2010), *enrichplot* (Yu, 2018), and *plotly* (Sievert, 2020).

## 3 Results

### Heat stress resolved the three pepper genotypes along distinct physiological trajectories

Physiological profiling revealed distinct temporal responses among the three *Capsicum annuum* genotypes (Figure 1). Because conductance was measured after a standardized 10-min equilibration period under laboratory conditions, these values describe genotype-dependent post-exposure responses under a common measurement protocol rather than steady-state conductance within the growth chamber. Representative plants illustrate phenotypic differences at selected stages of heat exposure (T1, T3, and T6; Figure 1A). The maximum efficiency of PSII in the light-adapted state (Fv′/Fm′) remained comparatively stable in GPC010350 throughout the time course, increasing slightly from 0.676 ± 0.032 at T0 to 0.705 ± 0.035 at T6, with mean values across the full time course ranging from approximately 0.68 to 0.73 (Figure 1B). By contrast, GPC014930 showed a progressive decline, from 0.684 ± 0.018 at T0 to 0.523 ± 0.081 at T6, whereas GPC003240 displayed a more moderate reduction, from 0.696 ± 0.022 at T0 to 0.636 ± 0.039 at T6. Differences among genotypes became most evident during the late phase of exposure: while Fv′/Fm′ was broadly comparable among genotypes at T0 (0.676–0.696), GPC010350 retained markedly higher values than GPC014930 and GPC003240 by T6. The effective quantum yield of PSII photochemistry (ΦPSII) showed a broadly comparable temporal pattern among genotypes and was therefore not included as a separate panel in the main figure. Complete PhotosynQ measurements, including ΦPSII and non-photochemical quenching (NPQt), are reported in Supplementary Table S4.

**Figure 1.**
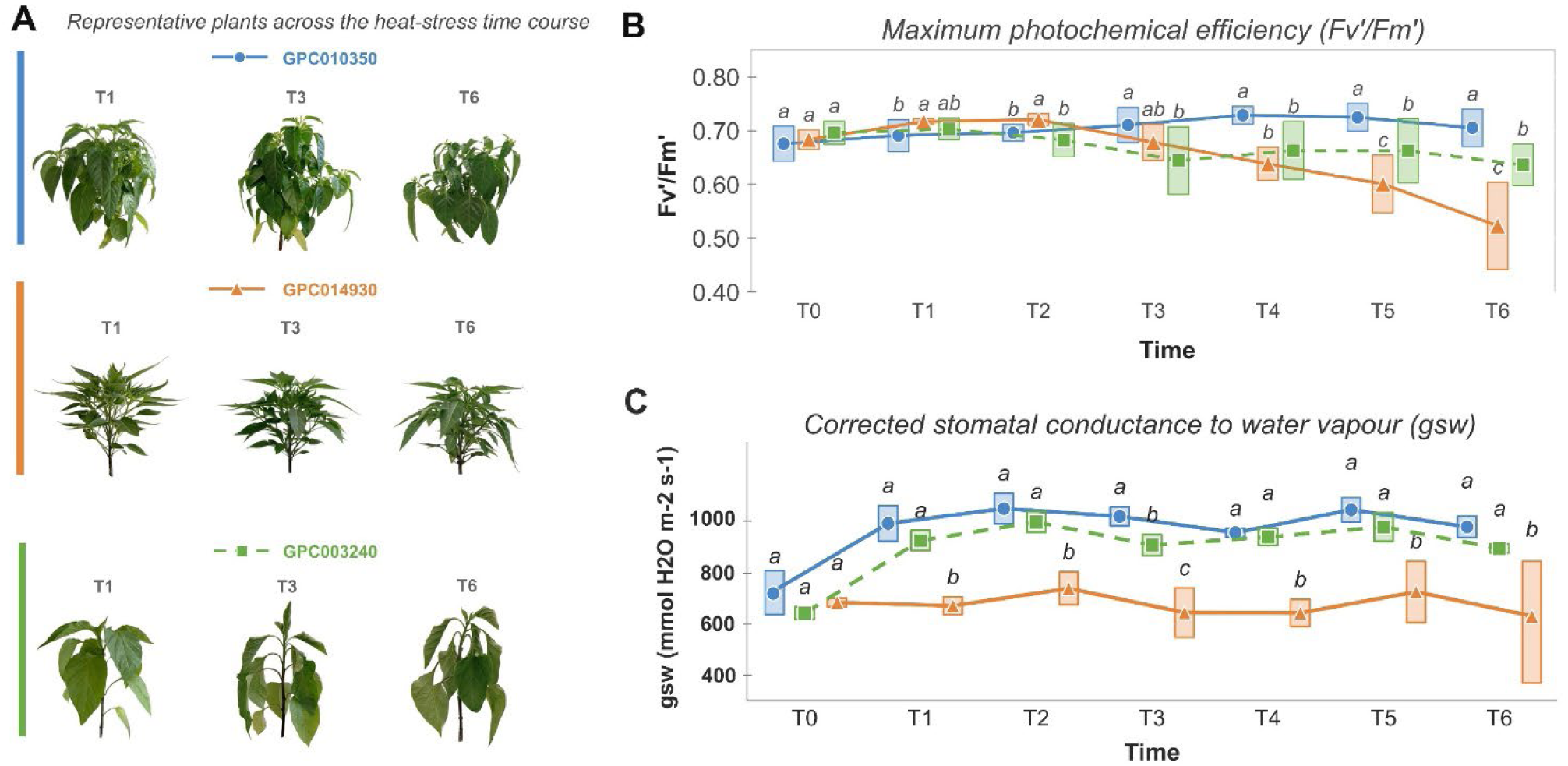
Phenotypic and physiological responses of three *Capsicum annuum* genotypes during prolonged heat exposure. (A) Representative plants at T1, T3, and T6. (B) Fv′/Fm′. (C) Psychrometrically corrected stomatal conductance to water vapour (gsw; sidedness = 1.0). Data are reported as means ± SD (n=3). Different letters indicate Holm-adjusted pairwise differences among genotypes within each time point (P < 0.05).

Psychrometrically corrected stomatal conductance to water vapour (gsw) also followed distinct temporal patterns among genotypes (Figure 1C). GPC010350 maintained comparatively high conductance throughout the time course, whereas GPC003240 increased after T0 and subsequently reached values similar to those of GPC010350. By contrast, GPC014930 generally showed lower conductance during the exposure period. The clearest separation emerged from the early-to-late phases of treatment, with GPC010350 and GPC003240 showing higher post-exposure conductance than GPC014930. Leaf water potential decreased between T0 and T6 across the three genotypes, but no significant genotype-dependent differences were detected (Supplementary Table S1; Figure S1).

Overall, physiological differentiation became more pronounced during prolonged exposure. GPC010350 combined comparatively stable photochemical efficiency with consistently high post-exposure stomatal conductance. GPC014930 showed progressive photochemical impairment and lower conductance, whereas GPC003240 combined increased conductance with a moderate late decline in Fv′/Fm′, defining a physiological trajectory distinct from those of the other two genotypes.

### Global transcriptomic profiling revealed major effects of genotype and heat exposure time

Principal component analysis of variance-stabilized expression values revealed a clear organization of the RNA-seq dataset, with PC1 and PC2 explaining 53.2% and 12.9% of the total variance, respectively (Figure 2A). Samples were distributed according to both genotype and exposure time, while biological replicates clustered closely within each genotype–time point combination, supporting the reproducibility of the dataset.

**Figure 2.**
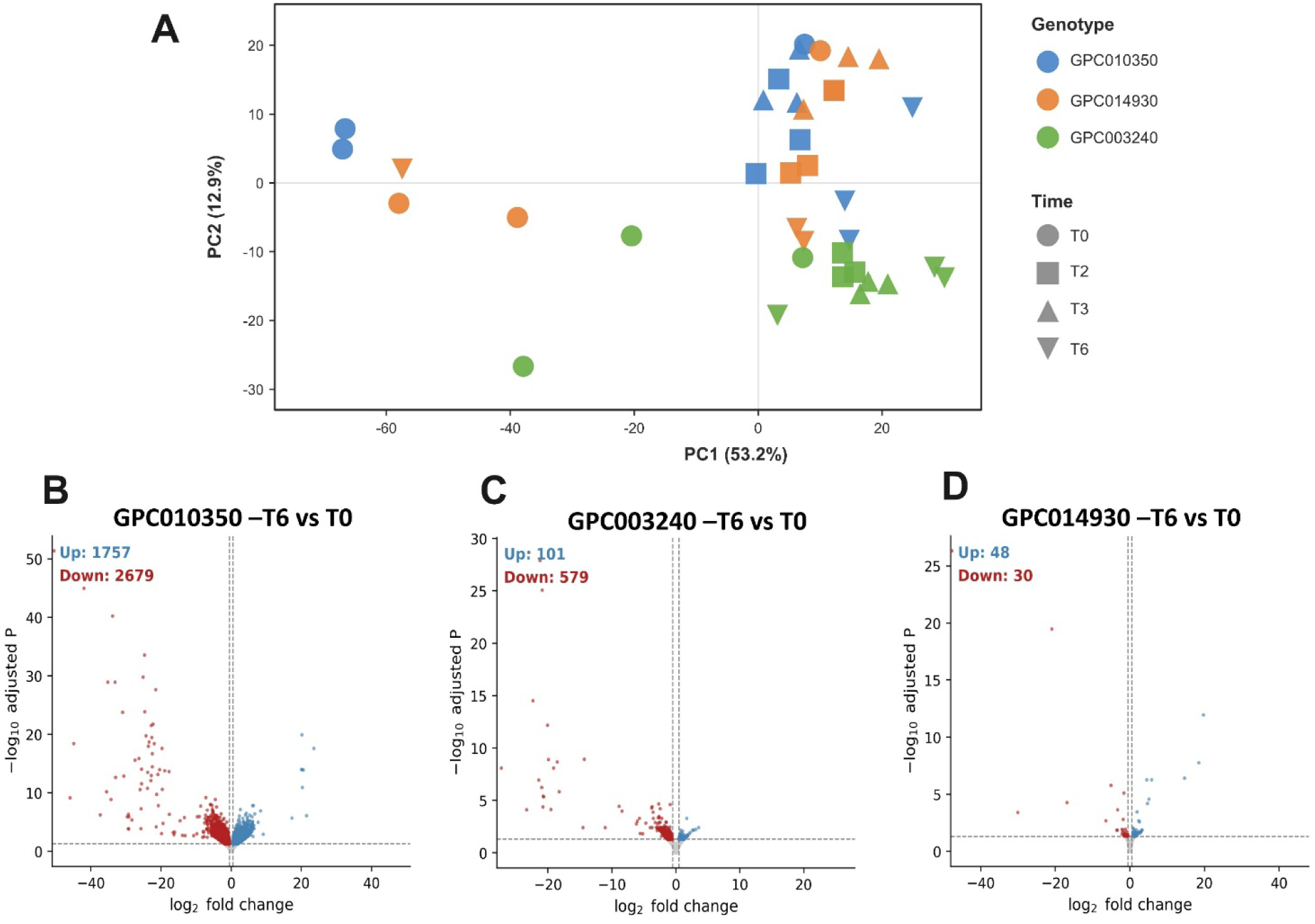
Transcriptomic responses during prolonged heat exposure. (A) PCA of variance-stabilized expression values, coloured by genotype and shaped by time point. (B–D) Volcano plots for T6 versus T0 in GPC010350, GPC003240 and GPC014930. Significant up- and downregulated genes are shown in blue and red, respectively; dashed lines indicate adjusted P < 0.05 and |log₂FC| > 1.

The T6-versus-T0 contrasts further revealed marked differences in the breadth of the late transcriptional response (Figure 2B–D). GPC010350 showed the largest response, with 4,436 differentially expressed genes, including 1,757 upregulated and 2,679 downregulated genes. GPC003240 displayed a more moderate response, with 680 DEGs, of which 101 were upregulated and 579 downregulated. By contrast, only 78 DEGs were detected in GPC014930 at T6, including 48 upregulated and 30 downregulated genes. Together, these results show that genotype and exposure duration jointly structured transcriptome-wide variation and that the three genotypes entered markedly different transcriptional states during prolonged heat exposure.

### Differential expression analysis reveals genotype-specific breadth and temporal dynamics of the transcriptome over the heat-exposure time course

Differentially expressed genes were defined using a BH-adjusted p-value < 0.05 and an absolute log2 fold-change > 0.5. Within each genotype, the dimension of the exposure-associated transcriptome varied markedly across the time course, revealing distinct temporal dynamics of transcriptional reprogramming (Figure 3).

**Figure 3.**
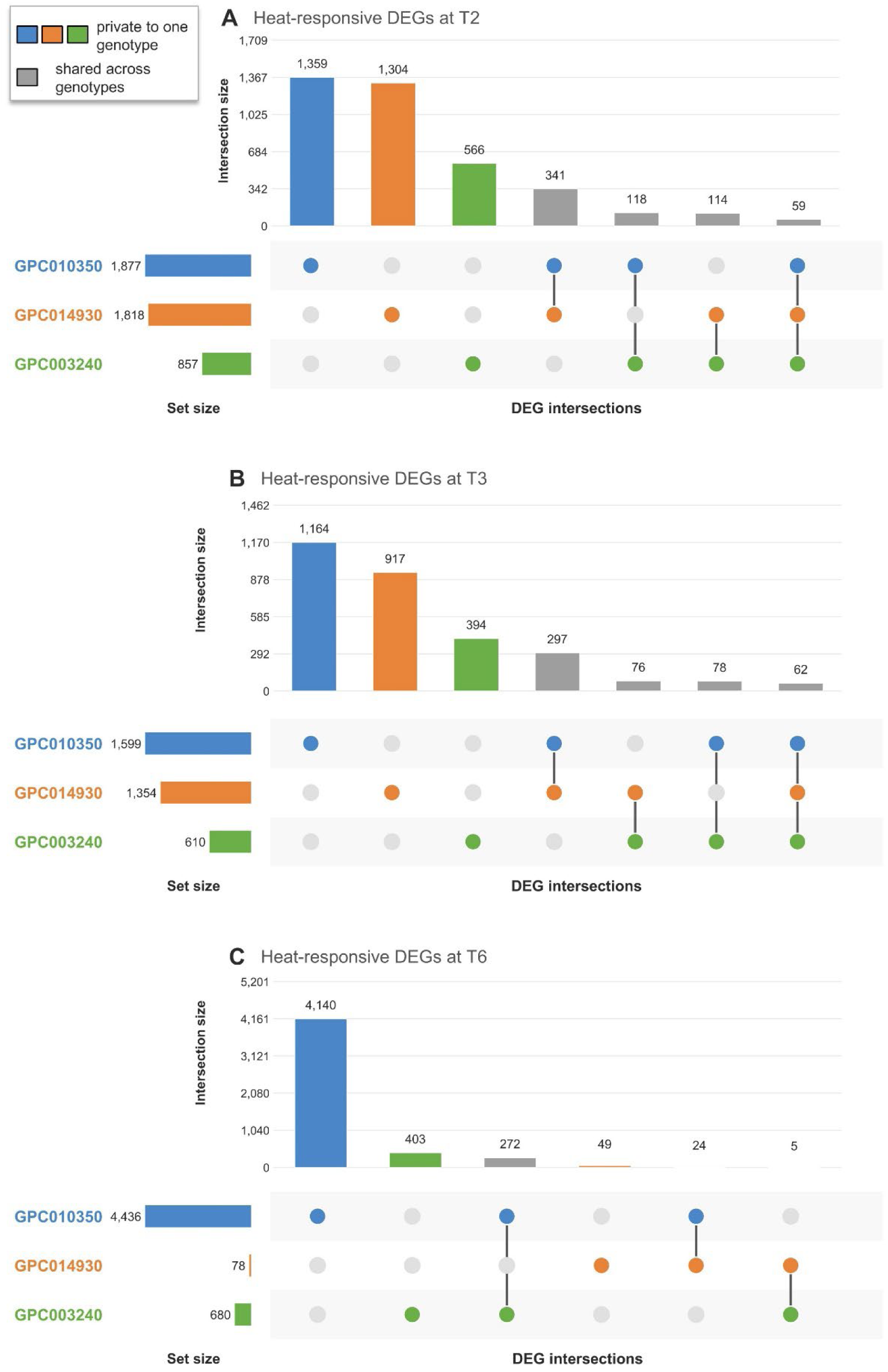
UpSet plots of exposure-associated DEG sets across genotypes at T2, T3, and T6. A) Exposure-associated DEGs at T2. B) Exposure-associated DEGs at T3. C) Exposure-associated DEGs at T6.

GPC003240 showed a moderate and relatively stable transcriptional response, with 857, 610, and 680 DEGs detected at T2, T3, and T6, respectively. GPC010350 displayed a pronounced late-expanding response: after 1,877 DEGs at T2 and 1,599 at T3, the number increased to 4,436 at T6, indicating extensive detectable transcriptional reorganization during prolonged exposure. By contrast, GPC014930 showed a broad early response, with 1,818 and 1,354 DEGs at T2 and T3, respectively, followed by a marked reduction at T6, when only 78 DEGs were identified (Table 1).

**Tabella 1.** Summary of differentially expressed genes (DEGs) per genotype and time point.

| Genotype | Time point | Total DEGs | Up | Down |
| --- | --- | --- | --- | --- |
| GPC003240 | T2 | 857 | 207 | 650 |
|  | T3 | 610 | 39 | 571 |
|  | T6 | 680 | 101 | 579 |
| GPC010350 | T2 | 1,877 | 602 | 1,275 |
|  | T3 | 1,599 | 356 | 1,243 |
|  | T6 | 4,436 | 1,757 | 2,679 |
| GPC014930 | T2 | 1,818 | 454 | 1,364 |
|  | T3 | 1,354 | 371 | 983 |
|  | T6 | 78 | 48 | 30 |

Comparison of genotype-specific DEG sets revealed both shared and genotype-restricted components of the exposure-associated transcriptome (Figure 3). At T2 and T3, 59 and 62 genes, respectively, met the DEG criteria in all three genotypes, representing only a small fraction of the total DEG union. No gene was shared across all three genotypes at T6. These patterns indicate limited overlap at the individual-gene level, even during the early response. However, absence from another genotype’s DEG set does not itself demonstrate a statistically significant difference between genotypes.

The largest pairwise overlap occurred between GPC010350 and GPC014930, which shared 341 DEGs at T2 and 300 at T3, indicating partial convergence of their early transcriptional responses despite their subsequent divergence. The genotype-restricted component also showed distinct temporal patterns. GPC003240 retained a moderate number of uniquely detected DEGs across the time course, with 575, 413, and 409 genes at T2, T3, and T6, respectively. In GPC010350, this component increased markedly at T6, from 1,367 and 1,170 genes at T2 and T3 to 4,161 genes at T6. By contrast, GPC014930 showed a reduction from 1,315 and 933 genotype-restricted DEGs at T2 and T3 to 50 at T6. Together, these descriptive patterns are consistent with increasing transcriptional divergence during prolonged exposure, although the interaction analyses provide the more direct evidence for genotype-dependent differences. Shared early-response genes, together with their annotations and genotype-specific fold changes, are reported in Supplementary Table S5.

### GSEA identifies shared early responses and late genotype-divergent functional programs

To characterize functional changes across the exposure time course, we performed Gene Set Enrichment Analysis (GSEA) of Gene Ontology (GO) terms using genes ranked by the DESeq2 Wald statistic and a *C. annuum* cv. ‘*Zhangshugang*’-specific GO annotation database. Because GSEA uses continuously ranked gene lists, it provides a complementary assessment that is independent of the fold-change threshold used for DEG classification. The main text focuses on the T6 within-genotype and genotype-by-time interaction contrasts in Biological Processes, where functional differences were most evident. T2 results and the Molecular Function and Cellular Component categories are reported in Supplementary Figure S3 and Supplementary Figure S4.

At T2, all three genotypes showed positive enrichment of processes related to RNA metabolism, ribosome biogenesis, translation, or organellar organization, indicating partial functional convergence during the early phase of exposure. In GPC003240, the strongest positive enrichments included miRNA processing (NES = 2.30) and ribosomal small-subunit assembly (NES = 2.29). GPC010350 showed enrichment of transcription by RNA polymerase III (NES = 2.20), mitochondrial RNA metabolic process (NES = 2.10), and regulatory ncRNA processing (NES = 1.91). In GPC014930, positively enriched terms included protein localization to chloroplast (NES = 2.26), chloroplast organization (NES = 1.73), and mitochondrion organization (NES = 1.76). Negative enrichment involved distinct signalling- and stress-associated processes, including jasmonic acid-mediated signalling and systemic acquired resistance in GPC003240, stomatal movement in GPC010350, and hormone-related signalling in GPC014930.

By T6, the functional profiles had become more distinct among genotypes (Figure 4A–C). GPC003240 showed positive enrichment of RNA splicing, large-subunit rRNA maturation, ribosome biogenesis, and rRNA metabolic processes, together with apoptotic process (Figure 4A). GPC010350 showed positive enrichment of amide biosynthetic process and DNA double-strand break repair through homologous recombination, whereas response to karrikin, floral organ development, and organic acid transport were negatively enriched (Figure 4B). GPC014930 showed negative enrichment of photosynthesis-associated processes, including light harvesting in photosystem I and plastid organization, as well as cell redox homeostasis (Figure 4C). DNA-repair-related gene sets remained positively enriched in GPC014930, whereas ribosome-biogenesis- and rRNA-processing-related terms were not prominent among its positively enriched processes.

**Figure 4.**
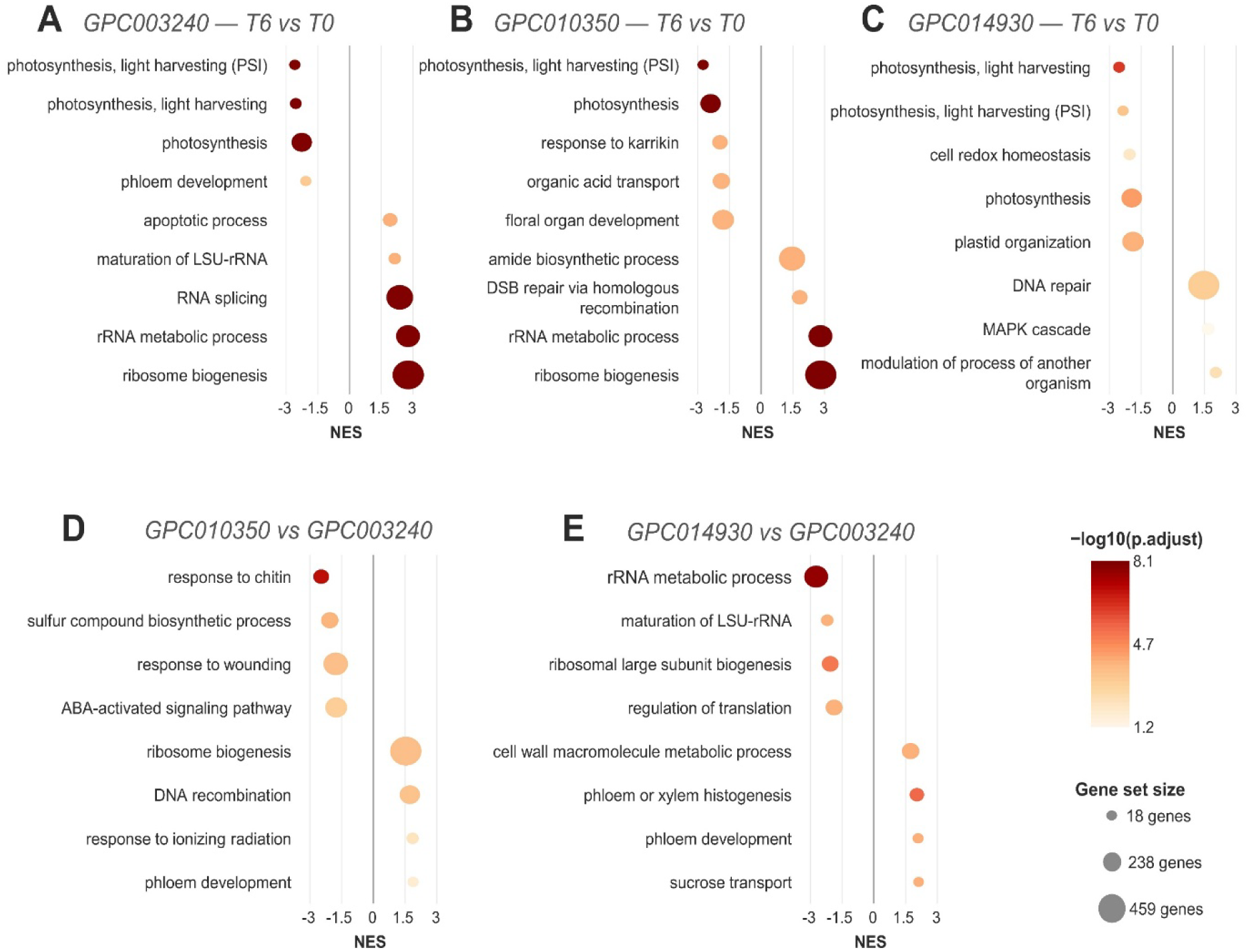
GSEA of GO Biological Process terms diverges among genotypes at T6. (A–C) Top activated/suppressed terms per genotype (T6 vs T0). (D–E) Genotype-by-time interaction terms relative to GPC003240. Dot size, gene set size; color, −log10(p.adjust).

GSEA of genotype-by-time interaction contrasts provided a direct assessment of differences in temporal responses relative to GPC003240 (Figure 4D,E). GPC010350 showed comparatively positive enrichment of ribosome biogenesis, DNA recombination, and response to ionizing radiation, together with comparatively negative enrichment of sulfur compound biosynthesis, ABA-mediated signalling, and wound response (Figure 4D). In GPC014930, comparatively positive enrichment involved phloem development, phloem and xylem histogenesis, sucrose transport, and cell-wall macromolecule metabolism. By contrast, rRNA metabolic process, large-subunit rRNA maturation, ribosomal large-subunit biogenesis, and regulation of translation showed comparatively negative enrichment (Figure 4E).

Overall, the GSEA results revealed partial convergence of RNA-, translation-, and organelle-associated functions during the early phase of exposure, followed by increasingly distinct functional profiles at T6. GPC010350 showed comparatively stronger enrichment of ribosome- and genome-maintenance-related processes, GPC003240 retained enrichment of RNA-processing and ribosome-biogenesis functions, and GPC014930 showed negative enrichment of photosynthesis-, plastid-, redox-, and translation-related processes.

### Global co-expression network analysis identifies exposure-time-correlated and genotype-associated modules

To identify co-expression programs associated with heat exposure and genotype identity at the systems level, we constructed a signed WGCNA network using all 36 samples. Genotype-dependent module trajectories were tested by fitting linear models to each module eigengene, with genotype, exposure time in hours, and their interaction as explanatory variables. Differences in T6-versus-T0 eigengene responses were additionally compared between genotype pairs. P-values were adjusted across modules using the Benjamini–Hochberg method. After retaining the 50% most variable genes and confirming sample quality, the analysis identified 29 co-expression modules, together with a grey module of unassigned genes (30 groups in total). The largest modules were turquoise (ME1), blue (ME2), brown (ME3), yellow (ME4), green (ME5), red (ME6), and black (ME7), comprising 5,295, 1,655, 983, 791, 628, 605, and 555 genes, respectively.

Correlation analysis between module eigengenes and heat exposure time identified co-expression programs associated with the temporal progression of stress (Supplementary Figure S5). After Benjamini–Hochberg correction for multiple testing across all 30 modules and four trait vectors, three modules remained significantly correlated with heat exposure time (Figure 5): ME12 (383 genes) showed the strongest positive correlation (r = 0.637, padj = 0.0012), followed by ME19 (248 genes; r = 0.498, padj = 0.035); both represent candidate co-expression programs sustained during prolonged heat exposure, whose eigengene values increased over time. ME5 (628 genes) was negatively correlated with time (r = -0.483, padj = 0.043), indicating a coordinated expression program that decreased during heat exposure. Additional modules (including ME1 and ME2) showed comparable nominal correlations with time but did not remain significant after correction and are not interpreted further.

**Figure 5.**
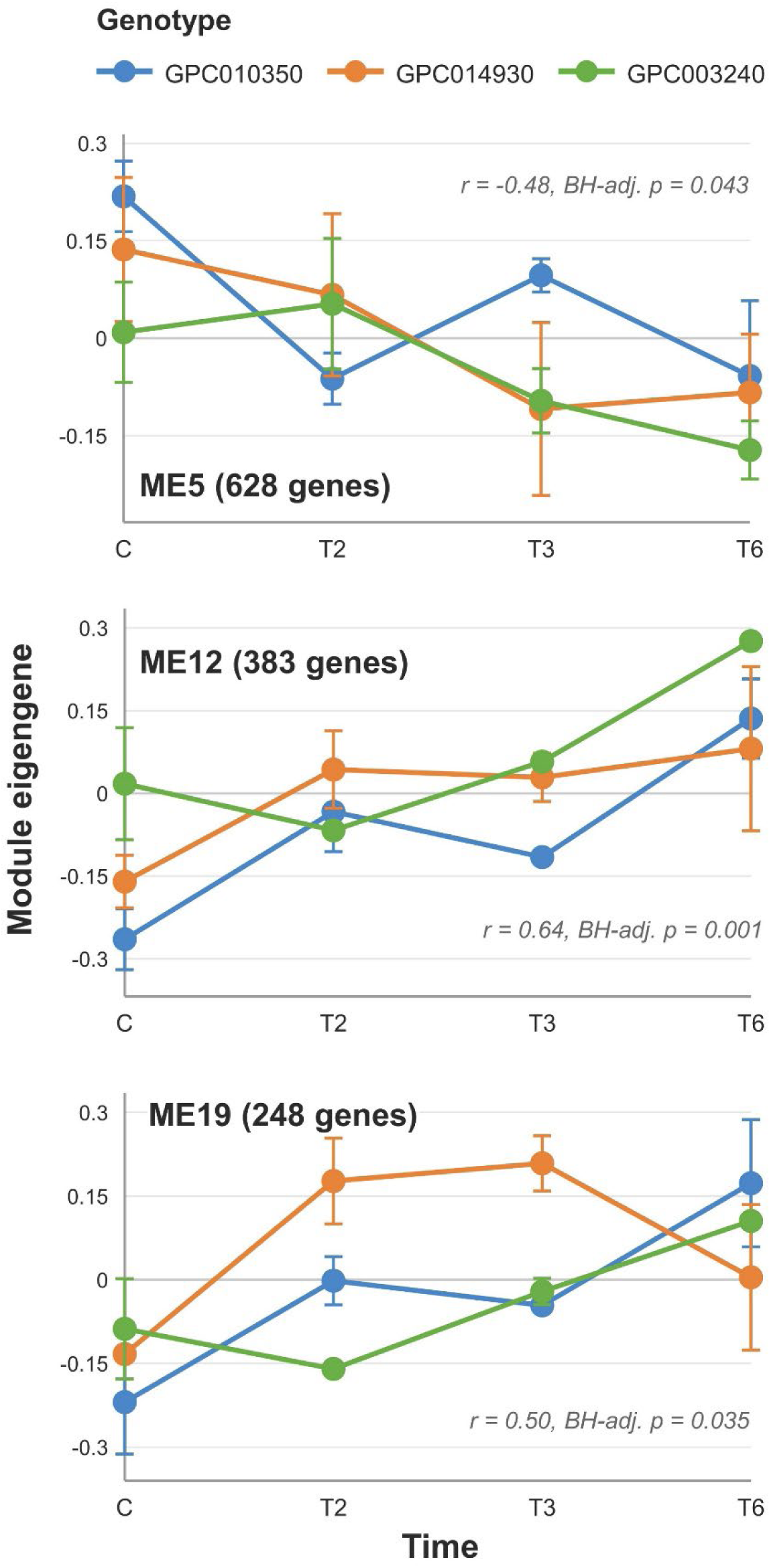
Time-responsive module eigengene trajectories in the global WGCNA network. Trajectories of the three modules significantly correlated with heat exposure time after Benjamini–Hochberg correction for multiple testing (mean ± SE) across GPC010350, GPC014930, and GPC003240.

Module–trait correlations also identified genotype-associated co-expression programs. After Benjamini–Hochberg correction, ME15 (318 genes; r = 0.780, padj = 1.2×10⁻⁶) and ME16 (299 genes; r = -0.846, padj = 9.9×10⁻⁹) were both robustly associated with GPC003240 identity, in opposite directions. Nominal trends suggested lower ME15 and higher ME16 eigengene values in GPC014930 and GPC010350 relative to GPC003240, but these individual genotype contrasts did not remain significant after correction (padj > 0.07 in all cases) and should be interpreted with caution. These modules capture a robust GPC003240-associated transcriptional background, though the present data do not provide independent statistical support for distinguishing GPC010350 from GPC014930 on this basis.

To formally test whether module trajectories differed between genotypes, we fitted a genotype × time model (module eigengene ∼ genotype × time) for each module and additionally compared T6-versus-T0 module responses between genotype pairs using linear models. After Benjamini–Hochberg correction, no module, including ME12, ME19, and ME5, showed a statistically significant genotype × time interaction, nor a significant genotype-specific difference in the T6 response in any pairwise comparison (all padj > 0.05). GPC010350 showed a statistically significant positive linear time trend in both ME12 (padj = 0.024) and ME19 (padj = 0.046). GPC003240 also showed a significant trend in ME12 (padj = 0.048) but not in ME19 (padj = 0.060), while GPC014930 showed no significant trend in either module (padj = 0.648 and 0.818, respectively). ME5 declined with comparable magnitude across all three genotypes, with no evidence of genotype-dependence. These patterns are therefore consistent with, but do not formally establish, genotype-dependent divergence at the co-expression module level; the DEG- and GSEA-based analyses (Sections 3.3–3.4) remain the primary evidence for genotype-dependent transcriptional responses. A full summary of all modules, including size, module–trait correlations, hub genes, and enriched GO terms, is provided in Supplementary Table S4.

### Prioritization of candidate genes

To characterize the biological content of the modules associated with exposure duration, WGCNA was integrated with Gene Ontology over-representation analysis and enrichment of module genes within DESeq2-derived DEG sets. The analysis focused on ME5, ME12, and ME19, the three modules significantly associated with exposure time. Candidate genes were prioritized by integrating module membership, significant differential expression in the relevant contrast, and functional annotation related to heat response, signaling, or cellular maintenance.

ME5, corresponding to the green module and negatively correlated with exposure duration, showed the strongest functional enrichment. After Benjamini–Hochberg correction, 41 Biological Process, 19 Molecular Function, and 45 Cellular Component terms remained significant, predominantly related to photosynthesis, light harvesting, photosystem I, chloroplast and thylakoid components, and chlorophyll binding. Highly connected genes included RBCS-1 (*Caz02g24250*), the *PSI* subunits *PsaD* (*Caz06g19780*) and *PsaE* (*Caz04g01890* and *Caz06g01410*), the PSII oxygen-evolving-complex proteins PsbO (*Caz02g06330*) and PsbP (*Caz07g10770*), and three light-harvesting complex proteins (*Caz03g07260*, *Caz04g24260*, and *Caz09g14760*). These nuclear genes encode chloroplast-targeted components of the photosynthetic apparatus and were significantly downregulated in GPC010350 at T2, with shrunken log₂ fold changes ranging from −2.07 to −2.74. ME5 genes were also over-represented among the T2 and T3 DEG sets of GPC010350 and GPC014930. ME5 therefore represents a broadly shared, negatively time-correlated photosynthesis-associated program rather than a response specific to one genotype.

ME12 and ME19, corresponding to the tan and lightyellow modules, respectively, were positively correlated with exposure duration, although no GO term remained significant after multiple-testing correction in either module. ME12 contained 383 genes, of which 226 were differentially expressed in GPC010350 at T6, corresponding to 59% of the module and a significant over-representation in the late GPC010350 DEG set. Highly connected genes included a *Pumilio*-family RNA-binding protein (*Caz04g05760*), Phloem Protein 2 (*Caz12g03980*), a MATE-family transporter (*Caz06g12750*), and glutamyl-tRNA reductase (*Caz01g31840*). All four genes were significantly upregulated in GPC010350 at T6. Their annotations suggest that ME12 integrates components of RNA regulation, transport, phloem-associated functions, and tetrapyrrole metabolism, although the absence of significant GO enrichment indicates that the module is functionally heterogeneous.

ME19 was enriched in the GPC010350 T6 DEG set and in genotype-dependent contrasts. *HSP101* (*Caz03g07770*), an *Hsp100/ClpB*-family chaperone, showed high module membership and significant induction in GPC010350 at T6. A heat shock transcription factor, *Caz03g27980*, was also significantly induced, although it showed more moderate module membership. ME19 additionally contained three genes annotated as dual-specificity phosphatases. *Caz05g20970* combined high module membership with significant induction in GPC010350 at T6, whereas *Caz05g21060* and *Caz05g21010* showed high module membership but did not meet the DEG criteria.

Overall, ME5 captured a shared decline in photosynthesis-associated expression, ME12 contained a large fraction of the late GPC010350-responsive transcriptome, and ME19 included established heat-response and signaling-related components. Integration of module membership, differential expression, and functional annotation prioritized *Caz03g27980*, *HSP101* (*Caz03g07770*), and *Caz05g20970* as candidates for further functional investigation. A complete summary of module composition, connectivity, enrichment, and candidate-gene annotations is provided in Supplementary Table S4.

## 4 Discussion

Integration of the physiological and transcriptomic datasets revealed genotype-dependent responses to prolonged heat exposure that became increasingly distinct over time. GPC010350 maintained comparatively stable photochemical performance and higher corrected stomatal conductance, whereas GPC014930 showed progressive photochemical impairment and a more limited stomatal response. GPC003240 followed a distinct but generally moderate trajectory. Physiological measurements were obtained after a standardized 10-min equilibration under common laboratory conditions and therefore reflect the combined influence of the preceding heat exposure and short-term adjustment to the measurement environment, rather than steady-state physiology inside the growth chamber at 40 °C. The physiological differences were accompanied by divergent transcriptional trajectories, indicating that genotypic variation may involve not only the magnitude of the early response but also its temporal organization during prolonged exposure.

Corrected stomatal conductance differed among genotypes across the time course, with GPC010350 and GPC003240 generally showing higher values than GPC014930 after the standardized equilibration period. Maintenance of relatively high stomatal conductance can contribute to heat avoidance when water is available by sustaining transpiration and facilitating leaf cooling. In Arabidopsis, elevated temperature promotes stomatal opening through a phototropin-dependent pathway involving activation of guard-cell plasma-membrane H⁺-ATPases, with 14-3-3 proteins contributing to stabilization of the active pumps; disruption of this response reduces transpiration and leaf cooling (Kostaki et al., 2020). In hot pepper, higher stomatal conductance and transpiration were associated with maintenance of photosynthetic activity in a comparatively heat-tolerant cultivar exposed to severe heat stress (Rajametov et al., 2021), and related genotype-dependent patterns have been reported in tomato (Wen et al., 2019). The higher values observed in GPC010350 and GPC003240 therefore cannot be considered direct evidence of greater transpiration or evaporative cooling during exposure at 40 °C but may indicate a greater capacity to maintain or rapidly re-establish stomatal opening after heat exposure. High post-exposure conductance coincided with comparatively stable Fv′/Fm′ in GPC010350, whereas GPC003240 showed a moderate late decline in Fv′/Fm′ despite similarly high conductance. Stomatal behavior may therefore have contributed to the physiological response but was insufficient to explain the differences in photochemical performance.

The temporal pattern of differential expression also differed markedly among genotypes. GPC010350 showed a pronounced expansion of detectable differential expression at T6, whereas GPC014930 displayed a broad early response followed by a substantial reduction in the number of genes meeting the DEG criteria at the same late time point. GPC003240 maintained a moderate and comparatively stable number of DEGs throughout the experiment. Thus, the magnitude of the early transcriptional response alone was not associated with the comparatively stable physiological profile of GPC010350. Instead, this genotype was characterized by continued and extensive transcriptional reorganization during prolonged exposure.

DEG numbers should nevertheless be interpreted cautiously. The number of genes passing a statistical threshold depends on effect size, dispersion, statistical power, and the selected fold-change and adjusted P-value cut-offs (Love et al., 2014). Moreover, significance in one genotype and non-significance in another does not itself demonstrate that their responses differ significantly; this requires a direct test of the genotype-by-time interaction (Gelman and Stern, 2006). DEG overlaps therefore provide a threshold-dependent description of temporal dynamics, whereas interaction contrasts and ranked enrichment analyses offer more direct evidence of genotype-dependent responses. The large late DEG set observed in GPC010350 is consequently best interpreted as extensive transcriptional reorganization rather than as evidence that a broader response is intrinsically more favorable.

Despite limited overlap among individual DEG sets, the early responses showed substantial convergence at the functional level. At T2, enriched processes were dominated by RNA processing, ribosome biogenesis, translation, ncRNA metabolism, and organellar organization. Heat stress affects multiple levels of gene-expression control, including RNA maturation, pre-mRNA splicing, rRNA processing, ribosome assembly, and translation (Kim et al., 2017; Darriere et al., 2022b; Dannfald et al., 2025). The shared enrichment of these functions may be therefore consistent with a common early reorganization of RNA and protein homeostasis. This shared functional component coexisted with genotype-dependent enrichment of signalling- and organelle-associated processes, suggesting that genetically distinct accessions may recruit partly different gene sets while affecting related cellular functions.

Genotype-dependent divergence became more pronounced at T6. In GPC010350, positive enrichment was retained for biosynthetic, RNA- and ribosome-associated, and genome-maintenance processes. DNA recombination and double-strand break repair were enriched in both within-genotype and interaction analyses. Elevated temperature can compromise genome integrity through ROS-mediated nucleotide damage, DNA strand breaks, and interference with replication and repair (Han et al., 2021). In Arabidopsis, the HSP90–HOS1–RECQ2 pathway has been linked to genome maintenance and acquired thermotolerance, providing experimental precedent for the involvement of DNA-repair mechanisms in heat responses (Han et al., 2020). The enrichment observed in GPC010350 is therefore consistent with the participation of genome-maintenance functions in its late response. Interaction analysis further indicated a comparatively stronger response of recombination- and genome-maintenance-related gene sets in GPC010350 than in GPC003240. These associations do not establish a direct contribution to the physiological differences, but they broaden the cellular processes associated with prolonged heat exposure beyond the canonical HSF–HSP response.

Co-expression analysis provided complementary information on the late response of GPC010350. ME12 contained a substantial proportion of genes differentially expressed in this genotype at T6 and included highly connected genes annotated as a Pumilio-family RNA-binding protein, a MATE-family transporter, Phloem Protein 2, and glutamyl-tRNA reductase. These annotations suggest possible contributions from post-transcriptional regulation, membrane transport, phloem-associated functions, and tetrapyrrole metabolism. Pumilio proteins can regulate RNA stability and translation, MATE transporters participate in the transport of diverse metabolites and signalling compounds, and glutamyl-tRNA reductase controls a key step in tetrapyrrole biosynthesis (Tam et al., 2010; Arae et al., 2019; Lu et al., 2019; Richter et al., 2019; Upadhyay et al., 2019). However, the absence of significant GO enrichment indicates that ME12 is functionally heterogeneous, and the roles of its individual genes during heat exposure remain to be determined.

ME19 showed a clearer connection with established heat-response components. The module contained HSP101 (*Caz03g07770*) and a heat shock transcription factor (HSF; *Caz03g27980*), both induced in GPC010350 at T6. HSFs regulate heat-responsive transcription, whereas *HSP101* is an *Hsp100/ClpB*-family chaperone involved in the recovery of heat-damaged proteins and acquired thermotolerance (Queitsch et al., 2000; Liu et al., 2011; McLoughlin et al., 2019). Their late induction is consistent with continued engagement of canonical heat-response components during prolonged exposure. ME19 also contained three genes annotated as dual-specificity phosphatases, including *Caz05g20970*, which combined high module membership with significant induction in GPC010350 at T6. MAPK phosphatases can modulate the duration and amplitude of environmental stress signalling (Ulm et al., 2002; Ding et al., 2018; Ayatollahi et al., 2022), making *Caz05g20970* a candidate for further investigation. Its specific function in pepper, however, cannot be inferred from annotation and co-expression alone.

The WGCNA results should be interpreted considering the statistical support for module-level differences. ME5, ME12, and ME19 were significantly associated with exposure duration in the global network, but genotype-by-time interactions for module eigengenes did not remain significant after multiple-testing correction. The module analyses therefore complement differential-expression and enrichment results by identifying coordinated gene sets and prioritizing candidate genes, but they do not independently establish genotype-specific regulatory programmes. Interaction-based differential-expression and GSEA analyses remain the primary evidence for genotype-dependent transcriptional divergence.

GPC014930 entered a contrasting late transcriptional state. Negative enrichment of photosynthesis, light harvesting, plastid organization, and cellular redox homeostasis accompanied its decline in photochemical performance. Photosynthetic electron transport, particularly PSII and its oxygen-evolving complex, is sensitive to elevated temperature, and disruption of excitation-energy transfer and electron transport can favor ROS production and oxidative damage (Allakhverdiev et al., 2008; Tang et al., 2022). The coordinated negative enrichment of photosynthetic and redox-related processes in GPC014930 is therefore consistent with broad alteration of functions related to energy capture and oxidative homeostasis. This interpretation remains associative: negative enrichment does not itself demonstrate loss of photosynthetic capacity or impaired control of oxidative stress. Interaction analysis also indicated a comparatively weaker response of ribosome-biogenesis- and translation-related gene sets in GPC014930 than in GPC003240, further distinguishing its late transcriptional profile.

ME5 captured the photosynthesis-associated response at the network level. The module was negatively correlated with exposure duration, enriched in chloroplast, thylakoid, light-harvesting, and photosystem functions, and contained highly connected nuclear genes encoding chloroplast-targeted components of the photosynthetic apparatus. However, ME5 genes were over-represented among early DEG sets in both GPC010350 and GPC014930, indicating that repression of photosynthesis-associated genes was not restricted to the genotype showing the strongest photochemical decline. Early downregulation of photosynthetic functions may therefore represent a broader component of the response to heat exposure, whereas later physiological divergence may depend on how this response occurred in combination with redox regulation, RNA and protein homeostasis, genome maintenance, and other stress-associated processes.

GPC003240 did not simply occupy an intermediate position in terms of DEG number. Its functional profile was characterized by sustained RNA-processing- and ribosome-biogenesis-related responses without either the pronounced late transcriptional expansion observed in GPC010350 or the combined photosynthetic, plastid, redox, and translation-related changes observed in GPC014930. The associations of ME15 and ME16 with GPC003240 further indicate that genotype-associated transcriptional backgrounds contributed to the global structure of the dataset. Because these modules primarily distinguished GPC003240 from the other genotypes, they did not independently resolve the contrast between GPC010350 and GPC014930. GPC003240 is therefore better described as showing a distinct, moderately responsive transcriptional profile than as representing a simple intermediate state.

Taken together, the results indicate that the three genotypes showed partially convergent early responses but increasingly distinct physiological and transcriptional profiles during prolonged heat exposure. In GPC010350, comparatively stable photochemical performance and higher corrected post-exposure stomatal conductance were associated with extensive late transcriptional reorganization, enrichment of RNA- and ribosome-associated and genome-maintenance functions, and induction of established heat-response components. In GPC014930, stronger photochemical impairment and lower post-exposure conductance were accompanied by negative enrichment of photosynthetic, plastid, redox, and translation-related functions. GPC003240 followed a distinct trajectory, combining increased post-exposure conductance with a moderate decline in photochemical performance and sustained RNA- and ribosome-associated responses. These observations suggest that genotype-dependent variation during prolonged exposure involved not only differences in the magnitude of the early transcriptional response but also differences in the temporal coordination of stress-response, maintenance, and metabolic processes. The conclusions should be considered within the scope of the experimental design. The study examined vegetative leaf responses under a single controlled heat regime and did not include a recovery phase. Moreover, RNA sequencing was performed at the common pre-treatment baseline and after two, three, and six days of heat exposure without time-matched control samples. Temporal transcriptional changes therefore cannot be completely separated from age- or time-dependent variation and are best interpreted as exposure-associated rather than as effects attributable exclusively to heat. Responses in reproductive tissues may also differ substantially from those observed in vegetative leaves, despite their direct relevance to pepper productivity (Erickson and Markhart, 2002; Reddy and Kakani, 2007; Gisbert-Mullor et al., 2023). Finally, differential expression, co-expression, and functional enrichment identify statistical associations rather than causal regulatory relationships. The genes and modules highlighted here therefore require validation through independent expression analyses, broader germplasm screening, and functional studies.

## 5 Conclusions

This study combined physiological profiling with time-course transcriptomic analyses to examine prolonged heat exposure in three *Capsicum annuum* genotypes. The genotypes showed partially convergent early responses but increasingly distinct profiles at later time points. GPC010350 maintained comparatively stable photochemical performance and showed extensive late transcriptional reorganization, including enrichment of RNA- and ribosome-associated functions, genome-maintenance processes, and established heat-response components. GPC014930 showed stronger photochemical impairment together with negative enrichment of photosynthetic, plastid, redox, and translation-related functions. GPC003240 displayed a distinct, moderately responsive profile.

Integration of differential expression, functional enrichment, and co-expression analyses identified ME12 and ME19 as components of the late GPC010350 response, whereas ME5 captured a broader photosynthesis-associated programme shared across genotypes. The heat shock transcription factor *Caz03g27980*, *HSP101* (*Caz03g07770*), and the dual-specificity phosphatase *Caz05g20970* represent candidates for subsequent functional validation through forward and reverse genetics. Overall, the results suggest that genotype-dependent variation during prolonged heat exposure involved not only differences in the magnitude of the early transcriptional response but also differences in its temporal organization. Within the limits of the experimental design, prolonged exposure was the phase in which genotype-dependent transcriptional trajectories and associated physiological profiles became most clearly distinguishable.

## Supporting information

Supplementary Figure S1. Leaf water potential across genotypes and time points.

Supplementary Figure S2 and Supplementary Figure S3. GSEA dot plots for (i) GO Molecular Function (S2) and Cellular Component (S3) terms for all eight

Supplementary Figure S2 and Supplementary Figure S3. GSEA dot plots for (i) GO Molecular Function (S2) and Cellular Component (S3) terms for all eight

upplementary Figure S4. Global WGCNA module-trait correlation heatmap. Heatmap of Pearson correlations between the 30 global module eigengenes and hea

Supplementary Figure S5. Per-genotype WGCNA module-trait heatmaps and module overlap summary.

## 7 Author Contributions

M.M: Conceptualization; Data Curation; Formal Analysis; Investigation; Methodology; Project administration; Resources; Software; Validation; Visualisation; Writing – original draft; Writing-review & editing. E.V: Data Curation; Formal Analysis; Writing – original draft; Writing-review & editing. F.S: Methodology; Visualisation; Writing – original draft; Writing-review & editing. A.M.M: Resources; Methodology; Writing – original draft; Writing-review & editing. L.B. Resources; Writing – original draft; Writing-review & editing. A.M.; Methodology; Visualisation; Writing – original draft; Writing-review & editing. A.A: Methodology; Visualisation; Writing – original draft; Writing-review & editing. C.C.: Visualisation; Supervision; Writing – original draft; Writing-review & editing. E.P: Investigation; Methodology; Funding acquisition; Project administration; Resources; Supervision; Validation; Visualization; Writing – original draft; Writing-review & editing.

## 6 Generative AI Statement

During the preparation of this manuscript, the author(s) used ChatGPT to improve the readability and language of the manuscript. After using this tool, the author(s) reviewed and edited the content as needed and take full responsibility for the content of the publication.

## 10 Supplementary Material

**Supplementary Table S1** – DESeq2 contrast definitions, coefficient specifications, and differential-expression summaries.

**Supplementary Table S2** – Sensitivity of DEG detection to alternative statistical and fold-change thresholds.

**Supplementary Table S3** – Summary of global WGCNA modules, module–trait associations, highly connected genes, and functional enrichment.

**Supplementary Table S4** – Physiological responses of three *Capsicum annuum* genotypes during prolonged heat exposure.

**Supplementary Table S5** – Shared exposure-associated core genes detected across the three genotypes at T2 and T3.

**Supplementary Figure S1.** Leaf water potential across genotypes and time points.

**Supplementary Figure S2 and Supplementary Figure S3.** GSEA dot plots for (i) GO Molecular Function (**S2**) and Cellular Component (**S3**) terms for all eight analyzed contrasts (three genotypes × T2/T6, plus the two interaction terms at T6).

**Supplementary Figure S4.** Global WGCNA module-trait correlation heatmap. Heatmap of Pearson correlations between the 30 global moduleeigengenes and heat-exposure time / genotype trait vectors, referenced in Section 3.5.

**Supplementary Figure S5.** Per-genotype WGCNA module-trait heatmaps and module overlap summary.

## Data Availability Statement

The raw sequencing reads generated in this study have been deposited in the NCBI Sequence Read Archive (SRA) under BioProject accession number SUB16378501.

