## Supplementary figures and images for "Genotype-dependent transcriptional trajectories during prolonged heat stress in *Capsicum annuum* L"

### Supplementary Figure S1. Leaf water potential across genotypes and time points.

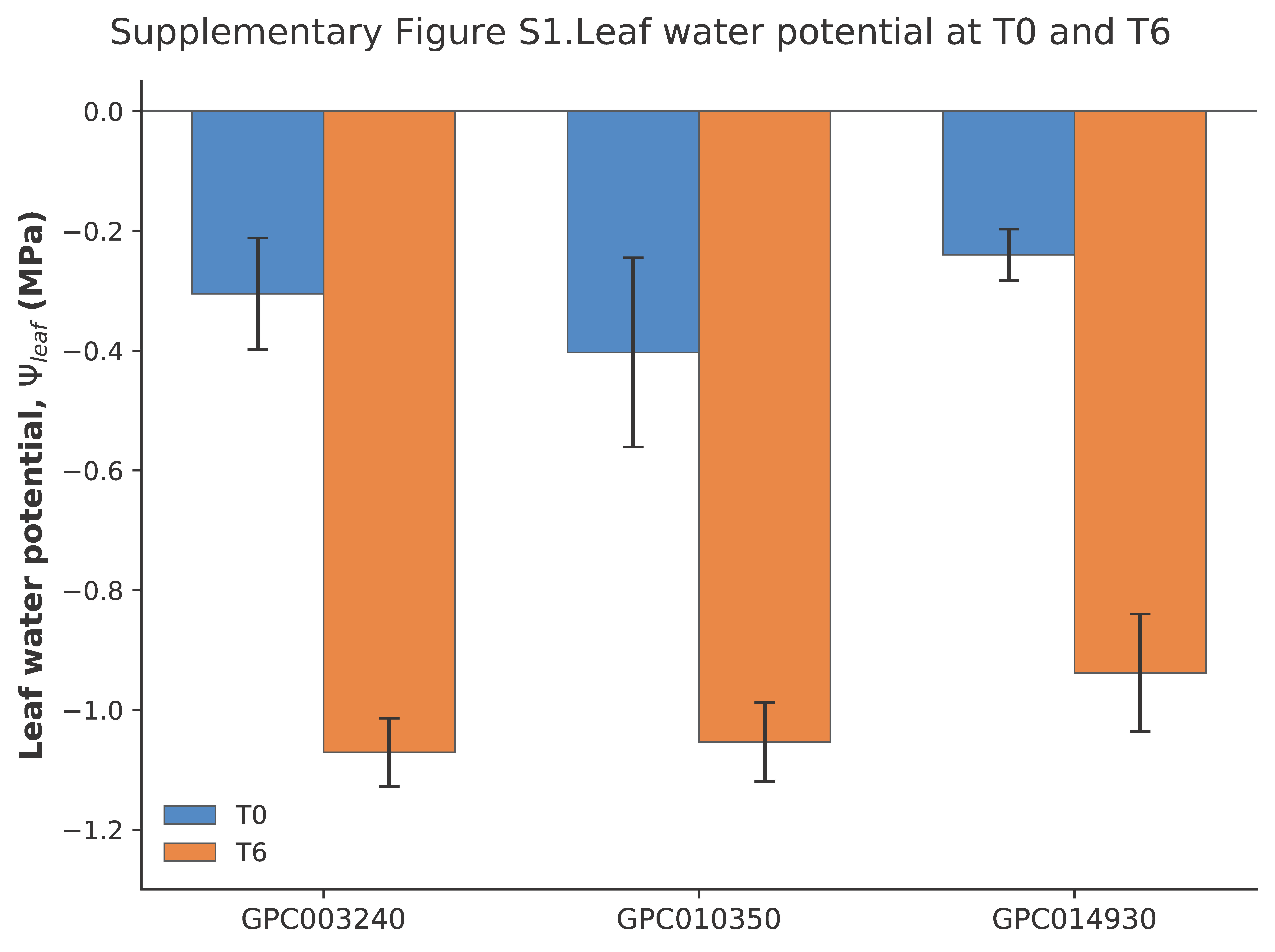

### Supplementary Figure S2 and Supplementary Figure S3. GSEA dot plots for (i) GO Molecular Function (S2) and Cellular Component (S3) terms for all eight

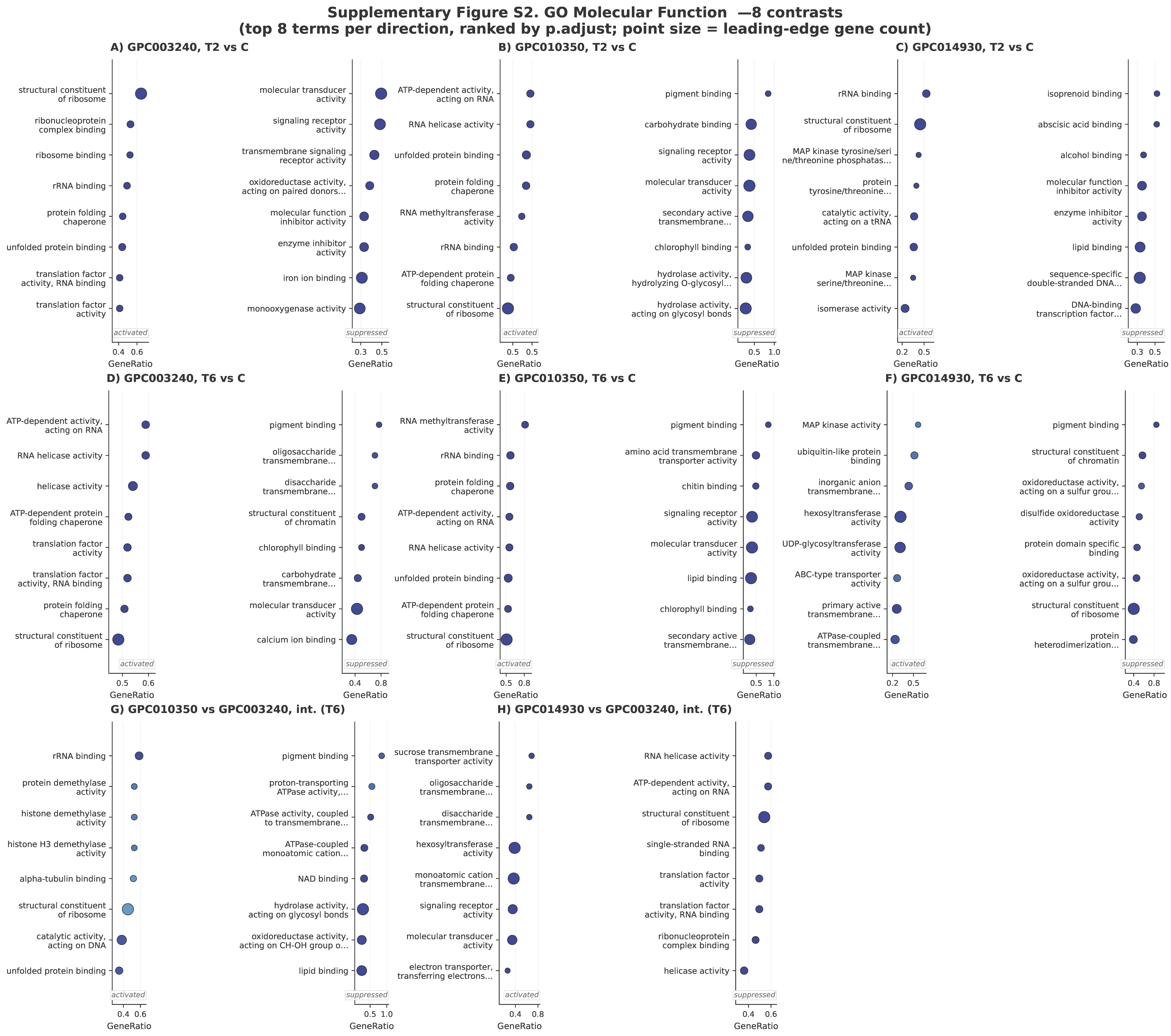

### Supplementary Figure S2 and Supplementary Figure S3. GSEA dot plots for (i) GO Molecular Function (S2) and Cellular Component (S3) terms for all eight

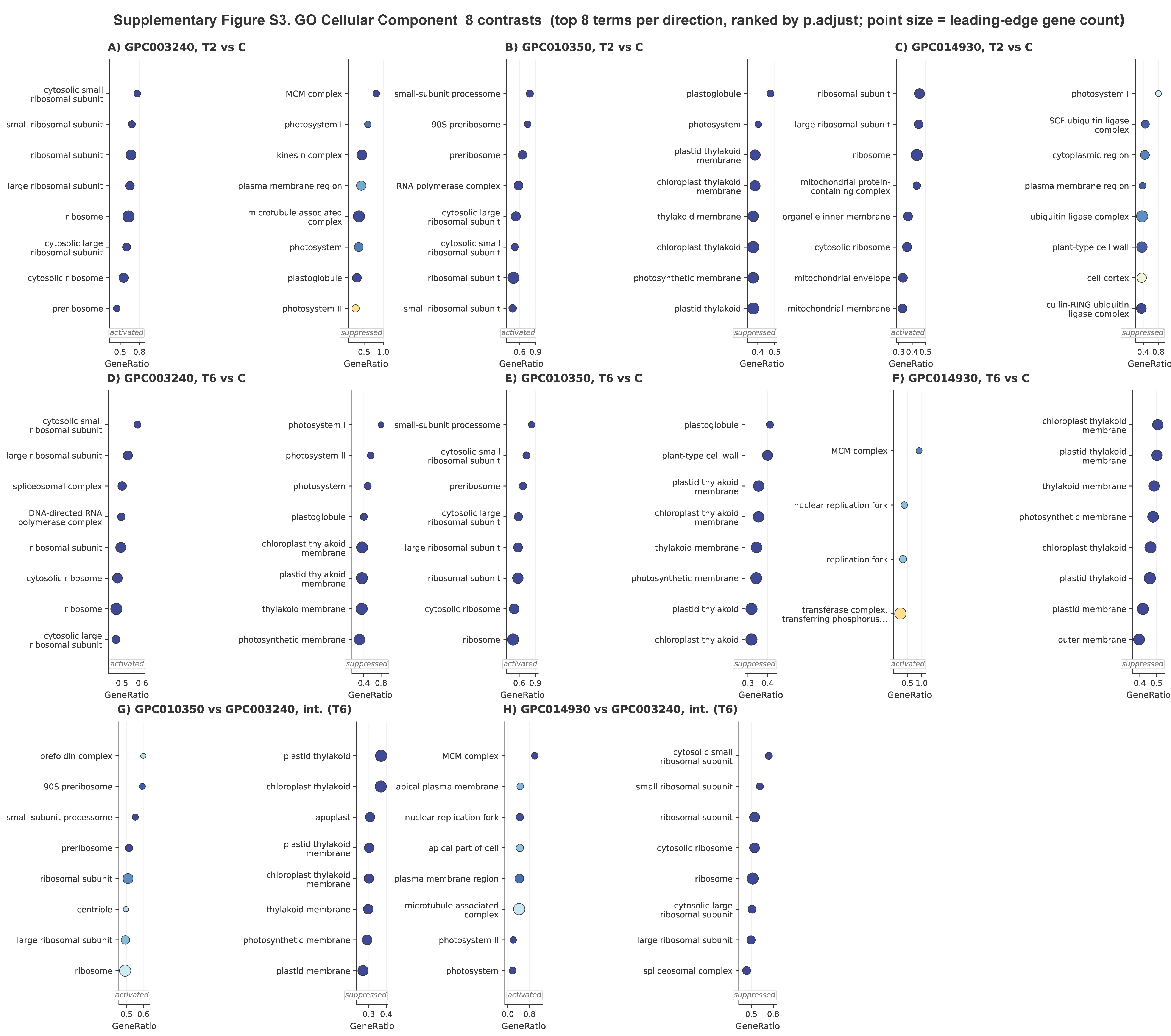

### Supplementary Figure S5. Per-genotype WGCNA module-trait heatmaps and module overlap summary.

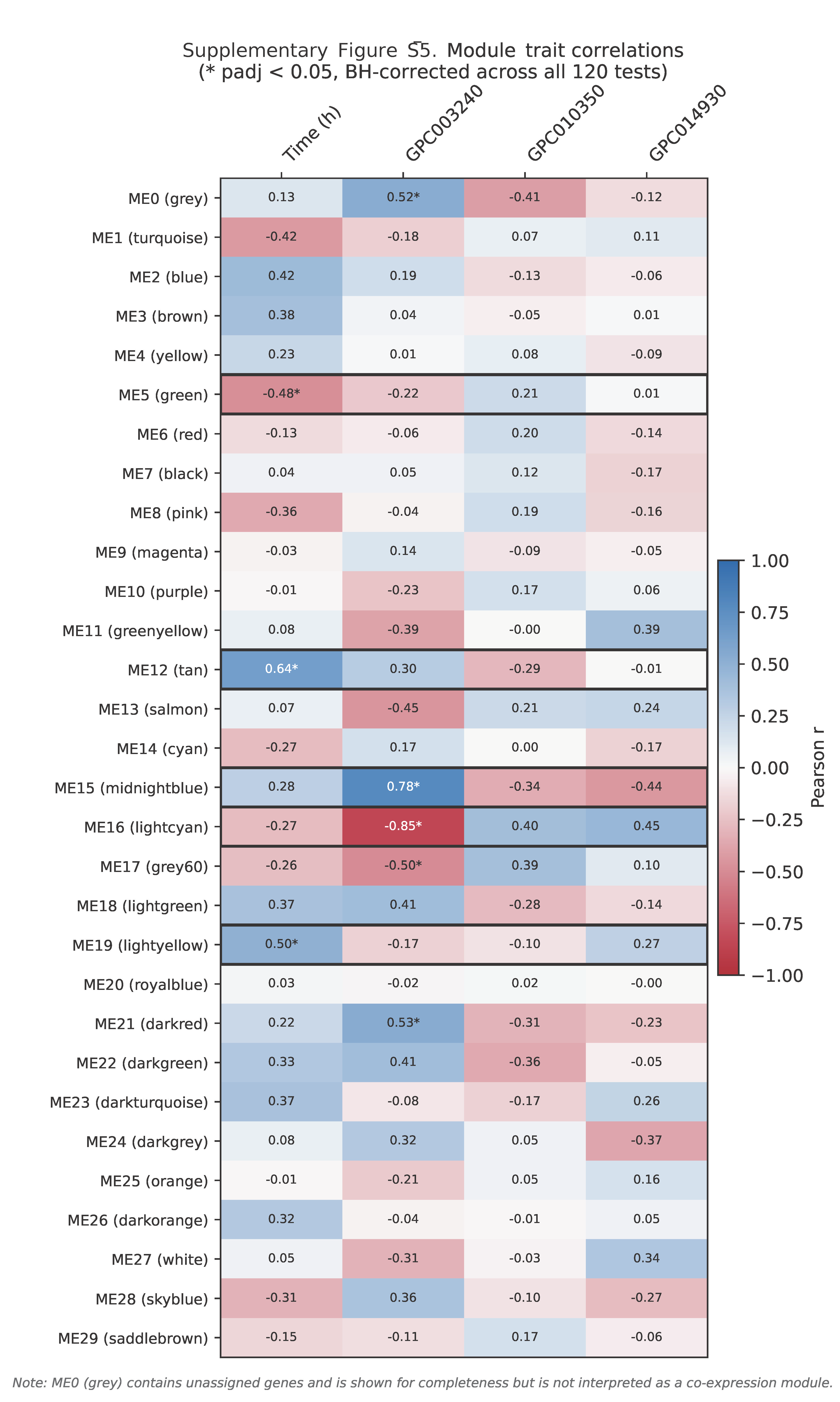

### upplementary Figure S4. Global WGCNA module-trait correlation heatmap. Heatmap of Pearson correlations between the 30 global module eigengenes and hea

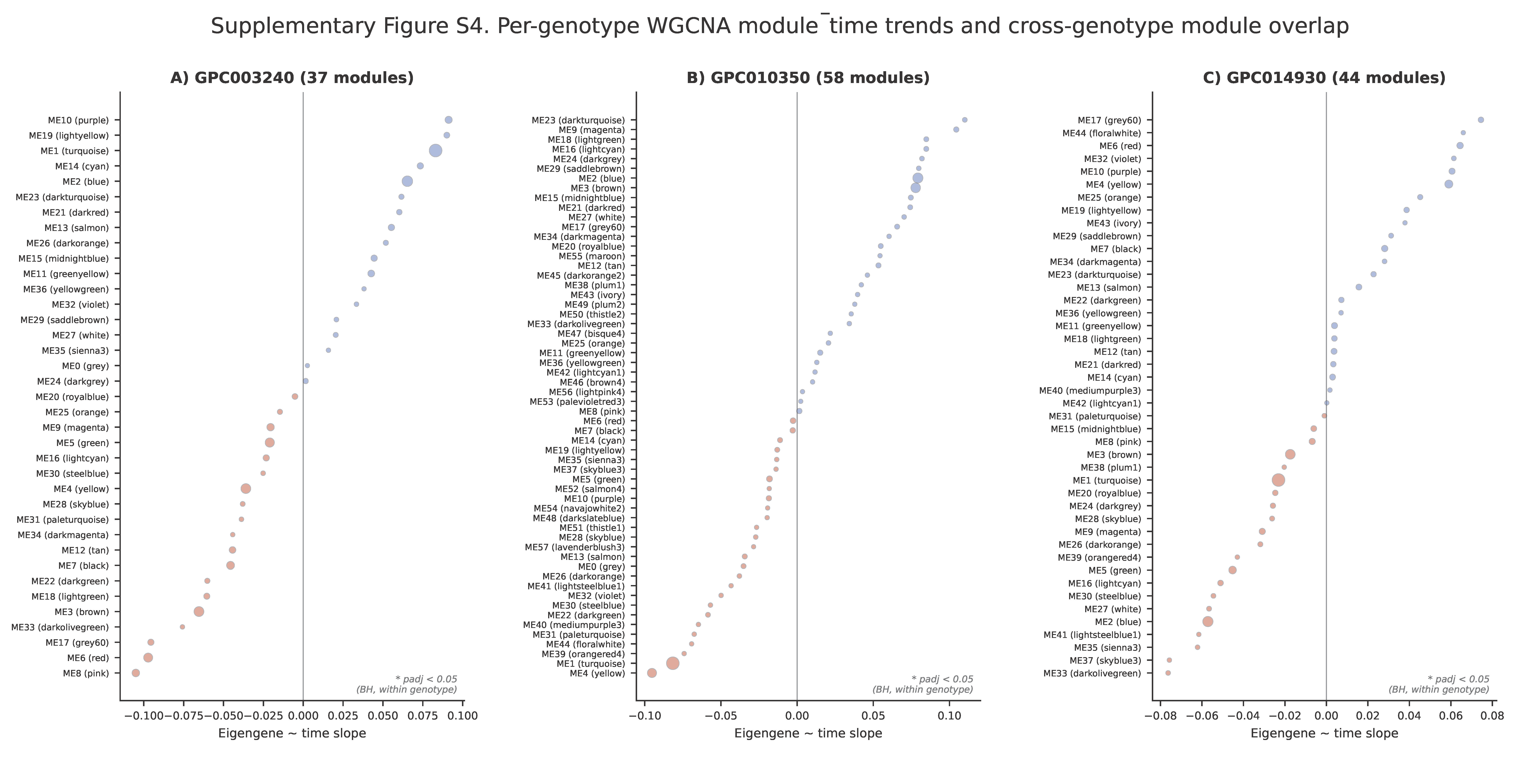
